# Genotype-environment association across precipitation regimens reveals the mechanism of plant adaptation to rainy environments

**DOI:** 10.1101/2024.06.21.600057

**Authors:** Simone Castellana, Paolo Maria Triozzi, Matteo Dell’Acqua, Elena Loreti, Pierdomenico Perata

## Abstract

In an era characterized by rapidly changing and less-predictable weather conditions fueled by the climate crisis, understanding the mechanisms underlying local adaptation in plants is of paramount importance for the conservation of species. As the frequency and intensity of extreme precipitation events increase, so are the flooding events resulting from soil water saturation. The deriving onset of hypoxic stress is one of the leading causes of crop damage and yield loss. By combining genomics and remote sensing data, today it is possible to probe natural plant populations that have evolved in different rainfall regimes and look for molecular adaptation to hypoxia. Here, using an environmental genome-wide association study (eGWAS) on 934 non-redundant georeferenced Arabidopsis ecotypes, we have identified functional variants for the gene *MED25 BINDING RING-H2 PROTEIN 1* (*MBR1*). This is a ubiquitin-protein ligase that regulates MEDIATOR25 (MED25), part of a multiprotein complex that interacts with transcription factors which act as key drivers of the hypoxic response in Arabidopsis, namely the RELATED TO AP2 proteins, RAP2.2 and RAP2.12. Through experimental validation, we show that natural variants of *MBR1* have a differential impact on the stability of MED25 and, in turn, on hypoxic tolerance. Our study also highlights the pivotal role of the MBR1/MED25 module in establishing a comprehensive hypoxic response. Our findings show that molecular candidates for plant environmental adaptation can be effectively mined from large datasets. This thus supports the need for the integration of forward and reverse genetics with robust molecular physiology validation of the outcomes.

## Introduction

Local adaptation in plant populations is the result of a combination of evolutionary history and phenotypic plasticity ^1,2^. Combinations of phenotypes that favor adaptation, resulting in increased fitness, ultimately depend on variations at the DNA level which are inherited and exchanged in the allele pool of natural populations ^3^. Understanding the genetic factors conferring local adaptation in an era characterized by rapidly shifting climate scenarios would enable plant biology to provide tools for the conservation and sustainable exploitation of biodiversity ^4,5^. The rise in global temperatures has an impact on the hydrological cycle ^6,7^. If on the one hand heatwaves and droughts are becoming increasingly common ^8,9^, on the other, there has been an intensification in extreme precipitation events, which lead to a higher occurrence of floods ^10,11^. Floods are among the most severe climate catastrophes in agriculture, with an estimated impact of 21 billion dollars for the decade 2008 – 2018 ^12^. Understanding the molecular basis of tolerance to abiotic stresses, including flooding, would break new ground in the development of new stress-resisting crop varieties ^13^.

During flood events, as heavy rainfall saturates the soil, plants may be partially submerged (a condition known as waterlogging) or completely covered by water (a state known as submergence). This condition inhibits gaseous exchanges that typically occur between the plant and the surrounding environment ^14–17^. As oxygen (O_2_) is a critical element for energy production and other vital metabolic processes, the hypoxic conditions that arise in flooded soils can severely impact plant growth and development ^18,19^. Plants have thus evolved sophisticated mechanisms to compensate for transient periods of oxygen deprivation. The primary strategy is to switch from oxidative phosphorylation, which requires oxygen, to anaerobic metabolism to generate the energy required ^20^. This process involves a finely tuned oxygen-dependent modulation of gene expression where one of the key pathways is the ethylene signaling pathway mediated by ERF-VII transcription factors (GROUP VII ETHYLENE RESPONSE FACTORS) ^21^. Under aerobic conditions, ERF-VIIs are constitutively oxidised by a class of plant enzymes named PLANT CYSTEINE OXIDASES (PCOs), which target the ERF-VII transcription factors to degradation via the proteasome ^22,23^, following the Cys branch of the N-degron pathway ^24,25^. Under hypoxia, ERF-VIIs escape degradation and are stabilized, enabling the transcription of core genes necessary for plant survival ^26^.

Despite comprehensive knowledge of oxygen-sensing mechanisms, forward genetic approaches such as genome-wide association studies (GWAS) could be exploited to reveal previously undiscovered regulatory elements in the hypoxic machinery. GWASs have helped identify new genetic variants associated with traits of interest and have complemented reverse genetic approaches focused on characterizing mutants and thus contributing significantly to our understanding of the genetic basis of plant adaptation ^27,28^.

Genetic variation is the primary driver of variation in a particular trait or phenotype, but GWASs involve a resource-intensive and time-consuming method of collecting phenotypic data. However, environmental factors can also have a significant impact on trait variation. In fact, the trait variations observed may not be due solely due to genetic factors. Instead, some of the variation may be due to environmental effects, leading to false positive or false negative results in the GWASs ^29^.

To overcome these limitations, environmental genome-wide association studies (eGWASs) have been used to identify associations between genetic diversity and pedoclimatic diversity in natural populations of plants and animals. These studies exploit the distribution of genetic diversity to infer past evolutionary processes and understand their contribution on extant variation ^30^. To date, eGWAS has been used on a wide range of different plant species, from tree species ^31,32^, to economically important crops ^33–35^. This has been made possible by the increasing availability of genomic information coupled with high-resolution, open-source climatic datasets deriving from remote sensing and re-analysis of climate data.

The model species *Arabidopsis thaliana*, due to its well-known genetics and wide global distribution, is an ideal candidate for eGWAS studies ^36,37^. Different studies have successfully used the eGWAS approach on Arabidopsis, revealing genes adapted to a wide range of environmental conditions; these include genes linked to arid climates ^38^, transcription factors involved in cold acclimatization ^39^, and sodium transporters that regulate salt tolerance based on proximity to the coast ^40^. Although eGWAS has a great potential in describing the molecular mechanism underlying adaptation, it is often held back by the lack of means to validate the importance of allele variation in contributing to the traits studied ^41^.

Based on these assumptions, we carried out an eGWAS to reveal potential candidate genes linked to areas of abundant rainfall. These genetic determinants can potentially shape an organism’s capacity to withstand the severity of rainfall-related challenges, such as waterlogging and submergence. We therefore used up-to-date databases regarding precipitation and soil characteristics, cross-referencing them with genetic data in a global collection of Arabidopsis.

We identified a strong association with single nucleotide polymorphisms (SNPs) present within the coding sequence of the *MED25 BINDING RING-H2 PROTEIN 1* (*MBR1)* gene. *MBR1* encodes for an E3-protein ubiquitin ligase that regulates the stability of a subunit of the Mediator complex ^42^, MEDIATOR25 (MED25), which in turn acts as a bridge between transcription factors and RNA polymerase II, ultimately regulating gene expression ^43^. MED25 interacts with two ERF-VII key transcription factors ^44,45^, which are key regulators of the hypoxic response in Arabidopsis.

In this study, we demonstrate how, through a cascade of events, the natural variants of *MBR1* impact the stability of MED25 and consequently the tolerance to hypoxic stress. We also demonstrate that the MBR1/MED25 module plays a central role in establishing a comprehensive hypoxic response. Our research underscores the efficacy of integrating eGWAS with experimental gene validation, which accelerates the identification of molecular factors that contribute to plant hypoxic adaptation. We believe that this approach opens new avenues for more effective conservation and harnessing of biodiversity aimed at strengthening resilience to the climate crisis.

## Results

### Climatic and Genetic Diversity in the collection

The 1001 genomes project (https://1001genomes.org/index.html) provides genomic sequencing data for 1135 Arabidopsis accessions collected in the wild. Starting with the overall dataset, we selected 934 nonredundant georeferenced ecotypes sampled across the global growth area of Arabidopsis (Supplemental Fig. 1a). We found that the genetic diversity existing in the data set was best summarized by 11 genetic clusters (Supplemental Table 1), in partially overlapping geographic regions (Fig. 1a; Supplemental Fig. 1c).

**Figure 1.**
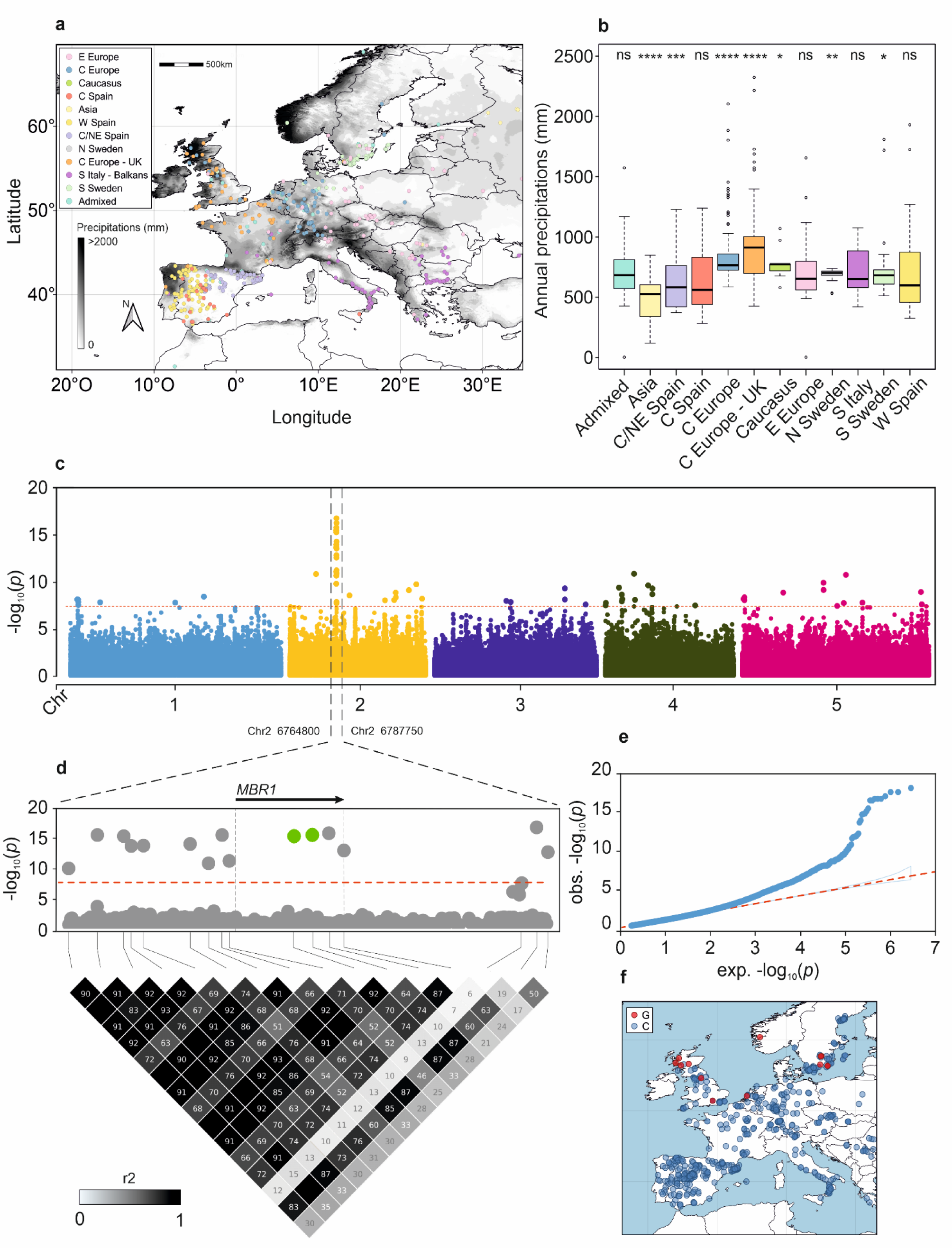
Association analysis with precipitation reveal variants in the *MBR1* gene. **(a)** Geographical location of the 11 clusters highlighted by ADMIXTURE; the grey color scale indicates the average annual rainfall expressed in millimeters for the period 1970 - 2020 **(b)** boxplot representing the distribution of accessions in relation to the annual rainfall expressed in mm of rainfall; statistically significant differences in the mean of all samples are indicated by asterisks (Student’s *t*-test; ns = non-significant; * = p < 0.05; ** = p < 0.01; *** = p < 0.001; **** = p < 0.0001). **(c)** Manhattan plot of eGWAS for the PC1 variable; the dashed red lines indicate the significance threshold set at *α* = 0.05 after Bonferroni correction. **(d)** Zoom on chromosome 2, the dashed vertical lines indicate the position of the *MBR1* gene; the green dots indicate the non-synonymous SNPs highlighted by the eGWAS analysis. On the bottom Linkage disequilibrium (LD) heatmap showing the LD patterns within the magnified region (**e)** Q-Q plot showing the observed versus expected -log10(*p*) values for the genome-wide association study for the PC1 variable. (**f**) Geographical distribution of the accessions carrying the most significant nonsynonymous SNP; the blue points represent the distribution of the reference allele (C/C), while the red points represent the distribution of the alternative allele (G/G)

According to the WorldClim dataset, rainfall in the collection ranged from 118mm to 2324mm. Most of the accessions were sampled in areas that experienced approximately 700mm / year (mean = 722,59mm; SD = 246,57). Gridded rainfall data deriving from different models may differ, so we compared WorldClim with other four datasets, finding high consistency (Pearson’s *r* from 0.73 to 0.99; Supplemental Fig. 2). We used a principal components analysis (PCA) to summarize precipitation data from different datasets in PC1, explaining 87.3% of the variance in rainfall data. This variable, representing the best approximation of precipitation variation in the area of study, was used in subsequent analyses. Together with this, correlations with various soil variables were analyzed, considering the close relationship between waterlogging phenomena and soil characteristics. An analysis was conducted to investigate the potential correlation between soil variables and precipitation variables, with the aim of confirming the independence of these datasets. The results demonstrated no correlation between the two sets of variables (Supplemental Fig. 3). This finding underscores that the soil and precipitation variables are entirely independent and do not share overlapping information. Together with this, we found that genetic clusters were characterized by different rainfall regimes (Fig. 1b, Supplemental Fig. 4), suggesting the possibility of local adaptation resulting from evolutionary processes.

### Environmental association analysis reveals an association with a gene located on chromosome 2

We conducted eGWAS on all the single rainfall dataset independently (Supplemental Fig. 5) as well as on the derived PC1. The resulting Manhattan plot for the PC1 is shown in Figure 1c. The complete list of the significant associations can be found in Supplemental Table 2. To identify a set of SNPs that are most likely to contribute to differences in protein-coding genes, we focused on polymorphisms occurring in the coding region of the gene, and specifically on missense variants. eGWAS revealed a highly significant peak located on chromosome 2, contributed by several SNPs in and around the gene *MBR1* (Fig. 1d). When performing linkage disequilibrium (LD) analysis in the locus, we found that all significant SNPs belong to the same LD block, meaning that it is not possible to resolve the association further (Fig. 1d). We found two SNPs with a minor allele frequency (MAF) of 0.02 in the coding region of the *MBR1* gene model; both these SNPs are predicted to cause missense mutations and have been chosen as prime candidates for further analyses. A list of accessions carrying the reference allele and the alternative allele can be found in Supplementary Table 3. We further explored environment associations conducting the eGWAS with month-specific precipitation data, finding higher significance for the *MBR1* association from October to January, the months characterized by higher rainfall values in their respective sampling locations (Fig. 2a - b). The same SNPs located in the *MBR1* gene resulted from an eGWAS considering the soil bulk density, a characteristic of the soil that is a proxy of soil water holding capacity (Supplemental Fig. 6a). This seasonal precipitation pattern aligns with the significant associations observed in the eGWAS, suggesting that the allelic variants of *MBR1* are likely adapted to environments characterized by higher winter rainfall.

**Figure 2.**
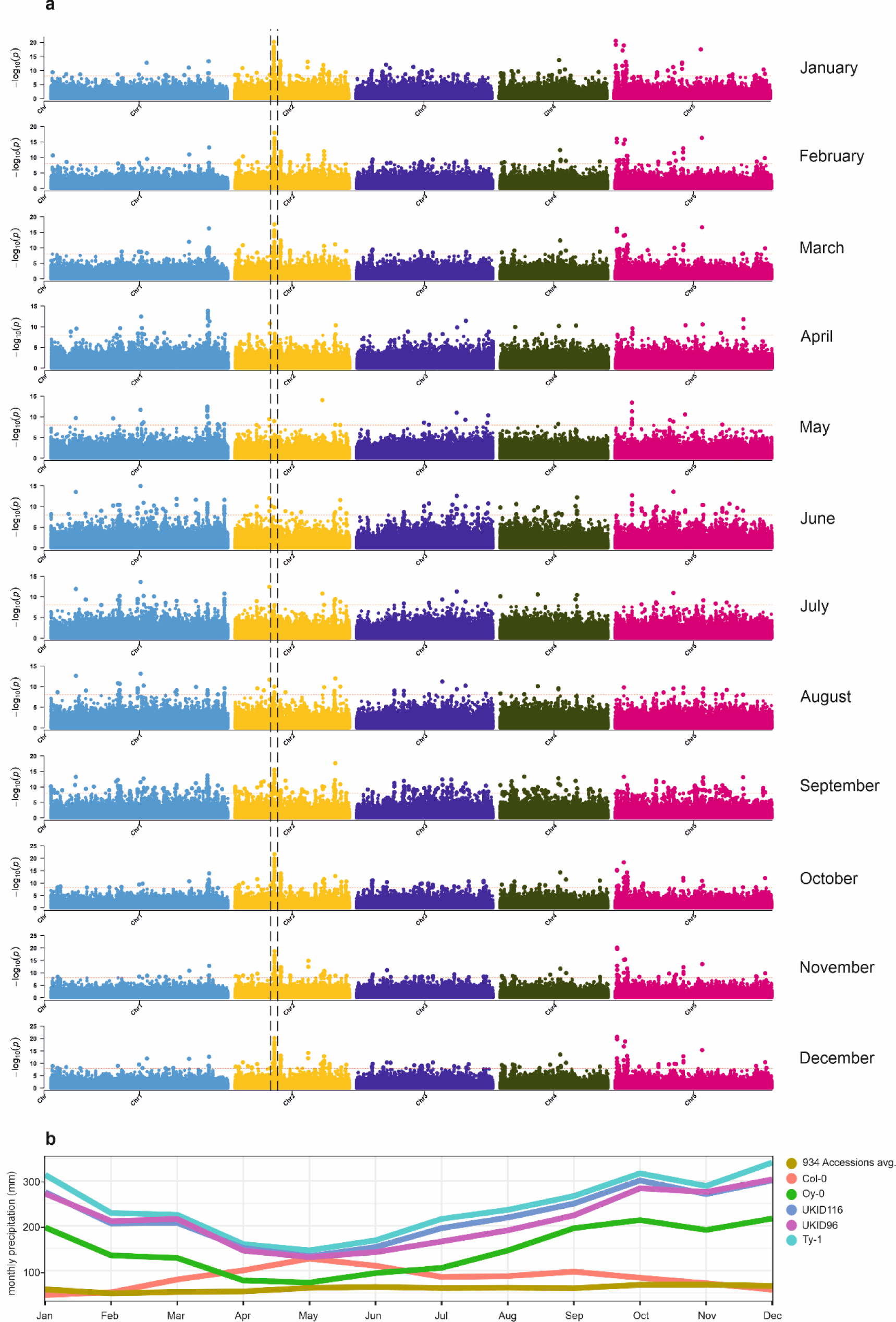
The marker-trait association tagging *MBR1* is more significant in months with higher rainfall. **(a)** Manhattan plots illustrating the eGWAS conducted with month-specific precipitation data for the period 1901 – 2020; dashed lines indicate the *MBR1* region as depicted in Fig. 1. **(b)** Monthly precipitation values (in mm) associated with accessions displaying alternative alleles at the *MBR1* locus. Col-0 and the average value of the entire set of 934 Arabidopsis show lower rainfall values during the boreal winter as compared with the ecotypes carrying the adaptive allele.

### Possible involvement of MBR1 in hypoxic tolerance

*MBR1* encodes a RING-type E3 ligase that acts as a regulator of MED25, a subunit of the Mediator complex, by directing its degradation in a RING-H2-dependent manner ^42^. MED25 physically interacts with transcriptional activators, including AP2/ERF, including two master regulators of the hypoxic response in plants, RAP2.2 and RAP2.12 ^44,45^. The close connection between *MBR1* and MED25, together with the interaction between MED25 and ERF-VII, led to the hypothesis of a possible link to the hypoxic response, where altered protein activity could affect the stability of MED25 and therefore the plant’s response to oxygen deprivation. To test the hypothesis, we first performed a submergence trial to assess the tolerance of the Col-0 genotype compared to natural ecotypes carrying the polymorphism in the gene, namely UKID96, UKID116, Ty-1, and Oy-0. The results indicated that all ecotypes showed a better performance compared to Col-0 (Fig. 3a). However, the ability of ecotypes to tolerate submergence better than the wild type could be related not only to the mutation in the *MBR1* gene, but also to a polygenic effect.

**Figure 3.**
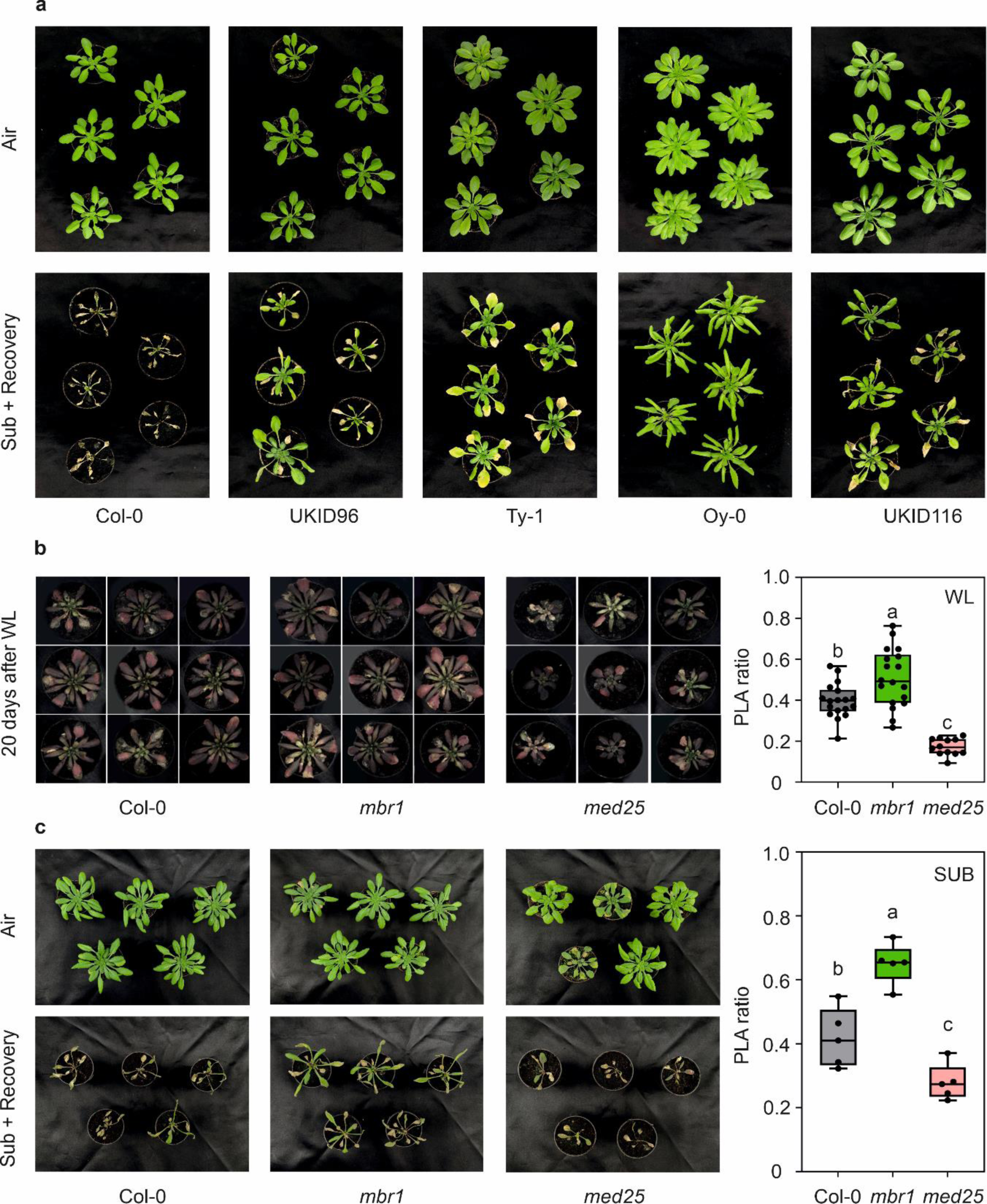
Submergence and waterlogging trials. **(a)** Effect of submergence on survival of Col-0 and accessions carrying the alternative version of *MBR1*. The plants were submerged for 48 h in the dark and transferred to normal 16h light / 8h dark conditions. Photographs were taken after 1 week of recovery. **(b)** Effect of waterlogging on the survival of Col-0, *mbr1,* and *med25* plants. The plants were kept in water for 20 days. Before phenotyping, the plants were randomised to help minimize potential bias and ensure that variations in environmental conditions were evenly distributed. Boxplot showing the PLA ratio assessed 20 days after the start of waterlogging. **(c)** Effect of submergence on the survival of Col-0, *mbr1,* and *med25* plants. Boxplot showing the PLA ratio assessed after 48h of recovery following 48h of submergence. In the boxplots, dots: represent single data points, whiskers denote the minimum/maximum values, the box defines the interquartile range, the centre represents the median, and box borders represent the lower and upper quartiles. Different letters (a, b, c) indicate differences in ANOVA tests (Tukey post hoc test, *P* < 0.05). The plant images were cropped and edited for clarification. The original unedited images for the waterlogging experiment can be found in Supplemental Figure 9.

To test whether *MBR1* is directly involved in the mechanisms of hypoxia tolerance, we employed both waterlogging and submersion experiments in our study. Initially, waterlogging was utilized for the Col-0, *mbr1* and *med25* knockout genotypes to simulate partial soil flooding, which is a common form of mild hypoxic stress encountered in natural environments (Fig 3b). This approach allowed us to observe variations in hypoxia tolerance under conditions that mirror real-world scenarios. To ensure consistency and validate our findings, we subsequently performed submersion experiments on Col-0, *mbr1* and *med25* genotypes (Fig 3c). The results, expressed as the ratio of Plant Leaf Area (PLA ratio) of waterlogged plants compared to those kept in the air, showed that *mbr1* tolerated the stress better than Col-0 in both experimental conditions, while *med25* showed the worst performance (Fig. 3b and c). These results indicate that the absence of *MBR1* results in a better hypoxic response, perhaps due to the greater stability of MED25. On the contrary, when MED25 is not present, the plant’s resistance to hypoxic stress is lower, providing additional evidence that MED25 plays a role in the plant’s hypoxic response.

### MBR1 acts as a repressor of the hypoxic transcriptional response

To further explore the role of *MBR1* in the hypoxic response, we evaluated gene expression levels using RT-qPCR. The expression levels of *MBR1* have been evaluated in Col-0 and the accession carrying the polymorphism after subjecting the plants to four hours of submergence. The results show a significant down-regulation of the MBR1 gene in all analyzed ecotypes (Fig. 4a). This repression of *MBR1* during submergence stress might be associated with a reduced activity of the protein, which could favor the stability of MED25 to ensure a more robust hypoxic response. Knockout *mbr1* plants, together with wild type, were subjected to submergence for 4 hours, with data collected during a time course (30 min, 1h, 2h, 4h), to check whether the absence of *MBR1* might play a role during hypoxic stress. We found that within the initial half-hour period, the expression levels for the core hypoxia genes *ADH1*, *PCO1*, *HRA1, PDC2, and LBD41* were up regulated compared to the wild type (Fig. 4b). This suggests that *MBR1* may normally dampen the hypoxia response in the wild type, possibly by affecting the stability of MED25. The gene expression levels were then equalised to the wild type after the first hour of submergence. This may be because in the wild-type *MBR1* is also down-regulated after the first hour of submergence, and eventually reaches similar expression levels to those of the *mbr1* mutant. The expression of *PGB1* was only slightly affected in *mbr1*, with upregulation 1h after submergence (Fig. 4b).

**Figure 4.**
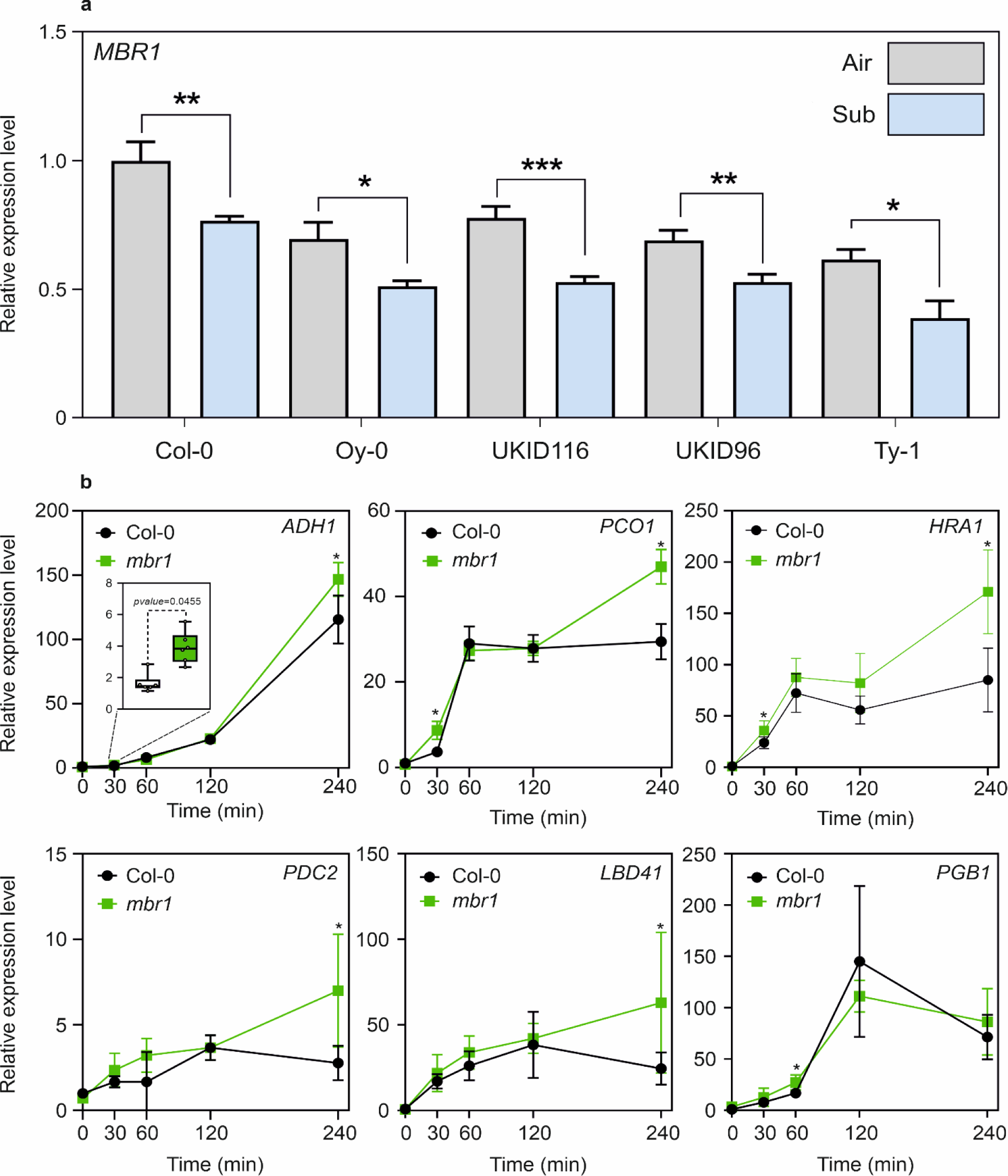
*MBR1* gene expression analysis and impact of the *mbr1* mutation on the expression of selected HRGs. **(a)** *MBR1* transcript level after 4h of submergence (Student’s *t*-test; ns = non-significant; * = p < 0.05; ** = p < 0.01; *** = p < 0.001; **** = p < 0.0001) **(b)** Time course of the expression level of a selection of anaerobic genes in submergence in Col-0 and *mbr1*; statistically significant differences are indicated by asterisks (Student’s *t*-test, unpaired comparison; * = p < 0.05). For all boxplots, the bottom and top of each box denote the first and third quartile, dots represent single data points, whiskers denote minimum/maximum values, the box defines the interquartile range, the center represents the median and box bounds represent the lower and upper quartiles.

### MED25 is more stable in ecotypes carrying the polymorphism

To test whether the MBR1 variant influences the stability of MED25, we performed a dual luciferase reporter assay to quantitatively measure the effect of *MBR1* on MED25 stability. Col-0 and ecotypes with the variant of *MBR1* were transformed with the *35S::MED25::FLuc* construct. Our findings showed that ecotypes with the MBR1 variant exhibited significantly higher relative luciferase activity compared to Col-0 (Fig. 5a).

**Figure 5.**
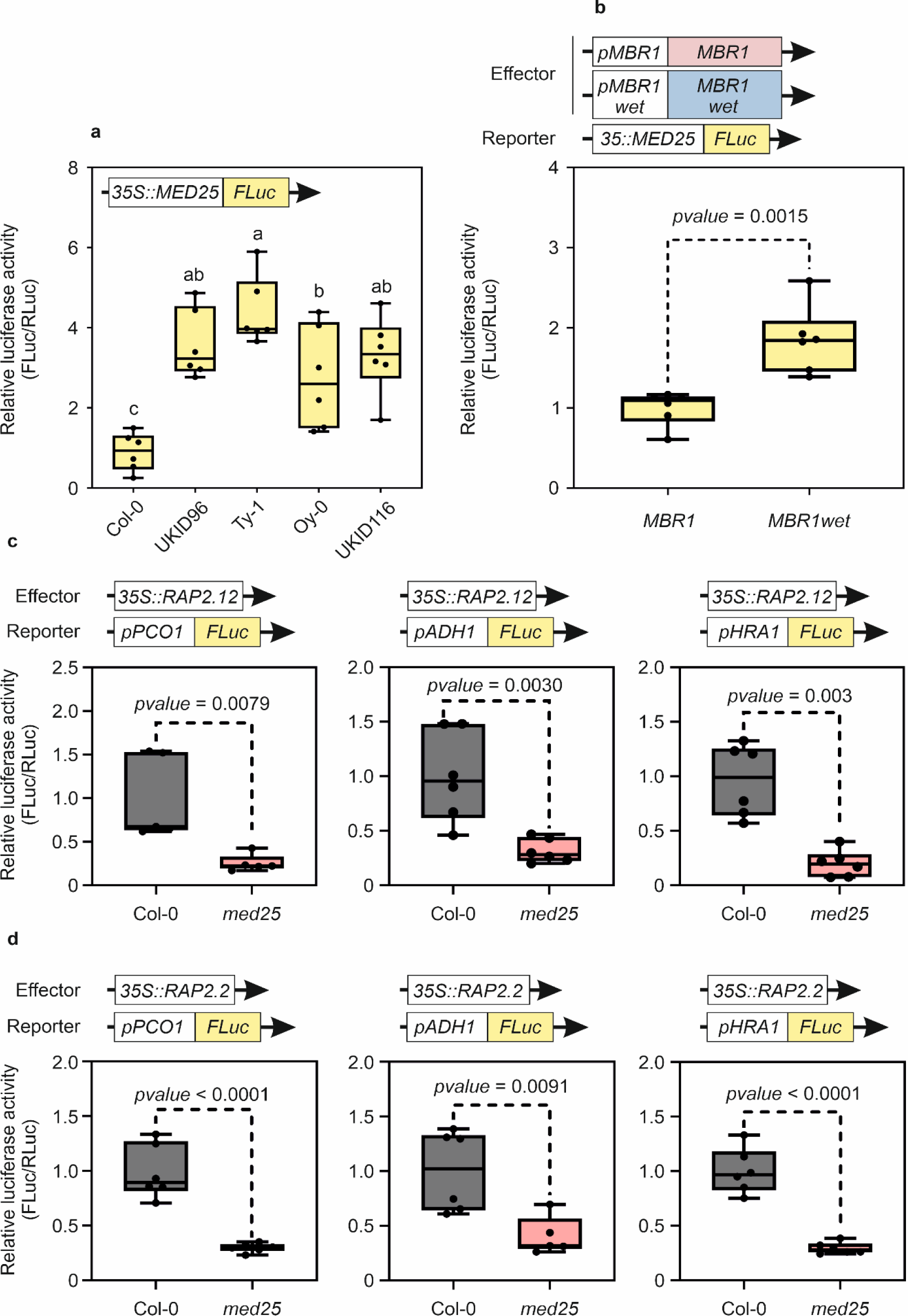
MED25 is required for the induction of HRGs by RAP2.12 and RAP2.2. **(a)** Relative 265 transcriptional activity of MED25 in the Col-0 protoplast and accessions carrying the variant of MBR1. **(b)** 266 Relative transcriptional activity of MED25 in protoplast of mbr1 transformed with the two versions of MBR1, 267 the wild-type version (-) and the one with the polymorphisms (+). **(c)** Relative transcriptional activity of the promoters of PCO1, ADH1 and HRA1 in protoplast of Col-0 and med25 transformed with an overexpressor of 269 RAP2.12 after 4h of hypoxia at 1%. (d) Same as (c), but protoplasts were transformed with the over-expressor 270 of RAP2.2. In the boxplots, dots represent single data points, whiskers denote the minimum/maximum values, 271 the box defines the interquartile range, the centre represents the median, and box borders represent the lower 272 and upper quartiles. Welch’s t test (two-sided) values are shown. Different letters (a, b, c, ab) indicate 273 differences in ANOVA tests (Tukey’s post hoc test, P < 0.05).

To validate whether the increased stability of MED25 was due to the alternative version of *MBR1* or rather to background variation, we cloned the wild-type *MBR1* version from Col- 0 and the *MBR1* version containing missense mutations from the Ty-1 ecotype as, according to the precipitation datasets, emerged as the accession that came from the rainiest environments. The vectors were then used for co-transformation starting from a knockout *mbr1* genotype to ensure that there was no endogenous form of the protein. The protoplasts isolated from the plants were co-transformed with one of the *MBR1* versions together with the *35S::MED25:FLuc* vector. The results showed that for the alternative version of *MBR1* there was greater MED25 activity, indicating that the stability of MED25 is related to *MBR1* and its alternative forms (Fig. 5b).

### MED25 is required for a complete hypoxic response

To validate the role of MED25 in the establishment of the hypoxic response, we implemented luciferase assays and protoplasts using Col-0 and *med25* knockout genotypes. Promoters of hypoxia-related genes, namely *ADH1*, *PCO1,* and *HRA1*, were cloned and fused with luciferase. These constructs were then co-transformed with effectors, represented by overexpressors of the *RAP2.12* and *RAP2.2* genes, into Col-0 and *med25* protoplasts. The results demonstrated a significantly reduced relative luciferase activity in all experiments (Fig. 5c-d). In the absence of *med25*, a full hypoxic response was unattainable, thus confirming the indispensable role of MED25.

## Discussion

The ability of a species to withstand the challenges of climate change hinges on its genetic diversity ^46^. The adaptation to shifting environments typically involves the gradual fine-tuning of molecular mechanisms, which occurs over evolutionary timescales. However, the current climate crisis is now fast-tracking these changes, putting pressure on organisms that are, likely, unable to adapt rapidly enough ^47^. The big data revolution in genomics and remote climate sensing has led to new ways of dealing with these challenges, leveraging diversity that is available in natural populations and connecting it to traits with clear adaptive significance. This knowledge could be used for the conservation and sustainable exploitation of biodiversity; however, it requires a clear description of the mechanisms underlying adaptation.

Here, we combine an eGWAS approach with molecular biology to identify and validate genes of adaptive significance, whose extant allelic variation is the result of evolutionary processes.

The results of environmental associations highlight the strong signs of association between rainfall and one gene, specifically *MBR1*. This gene is known to regulate MED25 stability^42^. MED25 is a subunit of the multiprotein complex and is involved in a wide range of plant functions ^43^. However, its key role is as a coactivator in the regulated transcription of genes dependent on RNA polymerase II. The Mediator complex functions as a bridge to convey information from gene-specific regulatory proteins to the basal RNA polymerase II transcription machinery. The mediator is recruited by promoters to direct interactions with regulatory proteins and serves as a scaffold for the assembly of a functional preinitiation complex with RNA polymerase II and general transcription factors ^48,49^. Among the transcription factors recruited by MED25, are *RAP2.2* and *RAP2.12* ^44,45^, which are two of the central regulators of the hypoxic response.

Recently, Schippers et al. ^50^ extensively validated the function of MED25 in hypoxic stress. Their study demonstrated that transcription factors belonging to the ERF family are recruited by MED25 to enable the full activation of hypoxia response mechanisms. Similarly, our findings reach the same conclusions, identifying MED25 as a crucial regulator of the plant’s hypoxic response. The absence of MED25 results in a significantly diminished induction of the core hypoxia-responsive genes. MED25 interacts with key ERFVII TFs, facilitating their binding to target gene promoters and thereby orchestrating a comprehensive transcriptional response to low oxygen conditions. Our data corroborate these insights, showing that without MED25, plants exhibit a notably weaker activation of genes essential for coping with hypoxic stress. This highlights MED25’s indispensable role in mediating the transcriptional network that underpins the plant’s survival and adaptation during episodes of oxygen deprivation.

The close connection between *MBR1*, MED25 and hypoxia led us to a compelling hypothesis. *MBR1* may have been subjected to natural selection in relation to rainfall, as it plays a role in modulating the hypoxic response, triggering a cascade of events that directly influence the stability of MED25, thus influencing the activity of ERF-VII transcription factors.

The influence of natural selection upstream of transcription factors is conceivable, as mutations within these factors could have detrimental consequences ^51^. Indeed, recent research has underscored the notable stability of the crucial regulatory factors that govern plant physiological responses, reducing the likelihood of mutations ^52^. However, one recent study revealed that accessions inhabiting distinct environments achieve adaptation through specific cis-element modifications, leading to increased constitutive expression of *RAP2.12*. This highlights how evolution can directly impact the regulatory elements involved in oxygen sensing ^53^. Alongside this, other studies have shown how specific mutations can contribute to the adaptation and thriving of Arabidopsis in particular environments. The study by Gou et al.^54^ identifies that natural variation in the *SVP* (SHORT VEGETATIVE PHASE) allele of the MADS-box gene in Arabidopsis thaliana has a pleiotropic effect on plant adaptation to contrasting environmental conditions. Specifically, they identified a non-functional mutation in the SVP-32V allele that alters normal regulatory interactions with target genes such as *GRF3*, *CYP707A1/3*, and *AtBG1*. This resulted in increased leaf size and greater tolerance to wet conditions but reduced drought tolerance. Similarly, the study by Kang et al.^56^ observed a specific insertion in the promoter region of the HPCA1 gene in the Tibet-0 ecotype, which increases gene expression and promotes adaptation to alpine environments.

The SNPs found within the *MBR1* gene has a low minor allele frequency (0.02) and the allele associated with high rainfall is restricted to populations in England and Scandinavia. Arabidopsis established relatively recently in northern Europe ^55,57^. Considering these findings, we may speculate that a correlation exists between the presence of the allele and the relatively recent northward expansion of Arabidopsis, that would be consistent with the species’ adaptive response to challenges posed by submergence. Certainly, this should be interpreted in the framework of a multi-locus process of adaptation, as corroborated by the correlation existing between genetic clusters and precipitation amounts (Supplemental Fig. 4).

Although geography and evolutionary history may have influenced the differentiation of these genetic groups, our results strongly imply that adaptive processes are associated with precipitation patterns. Together with this evidence, we found that *MBR1* was significantly associated with soil bulk density, which bears no correlation with rainfall, further strengthening the possible role of the gene in the response to hypoxia during waterlogging.

The close correlation between the bulk density and the possibility of soil flooding is interesting. Soils with a high bulk density are typically more compacted and have a limited capacity to absorb and drain water during extreme rainfall events. As a result, water may accumulate on the surface, leading to poor drainage and increasing the risk of waterlogging and therefore the establishment of a hypoxic environment ^58^.

One frequent significant limitation of the current eGWAS literature is the lack of experimental validation for genes identified through association studies ^41^. Our aim was to specifically address this limitation by conducting comprehensive experimental investigations to confirm the role of the candidate genes identified. This approach aimed to bridge the gap between genetic associations and functional insights, thereby contributing to a more comprehensive understanding of the mechanisms underpinning plant adaptation.

Through experimental investigations, we successfully revealed the role of *MBR1* in hypoxia stress. First, we observed that plants from rainy environments displayed superior tolerance to submergence stress. However, since this resistance could be putatively linked to a multigenic effect, we implemented *MBR1* and *MED25* knockout lines in waterlogging and submergence experiments, revealing that in both conditions *mbr1* provided better resistance than Col-0 and *med25* (Figure 3). This clarified the potential involvement of the MBR1/MED25 module in the hypoxia response machinery, further supported by qPCR experiments demonstrating superior hypoxic gene induction in *mbr1* compared to Col-0 (Figure 4). Luciferase assays indicated that accessions from rainy environments exhibited greater stability of MED25, due to reduced *MBR1* activity (Figure 5b). Lastly, we provide the first direct evidence of the essential role of MED25 in establishing a comprehensive hypoxic response (Figure 5c-d).

Our research is aimed at understanding the intricate ways plants adapt to their surroundings. Our results show existing databases, such as the 1001 Genomes Project, already hold a treasure of information waiting to be extracted. We demonstrate that tools such as eGWAS can serve as a compass for unearthing genes closely intertwined with species adaptation to distinct, often harsh environments. The discovery of a mutation in the *MBR1* gene, which leads to a cascade effect on hypoxia tolerance, offers a fascinating glimpse into the intricate molecular mechanisms underlying plant adaptation to challenging environments.

Exploring how organisms perceive oxygen and respond to extreme environments ^59^ not only enriches our knowledge but also charts promising pathways for comprehending adaptive strategies amid evolving climate challenges. In addition to unveiling a novel function of the *MBR1* gene, we have ventured into uncharted territory by revealing the role of MED25 in the hypoxic response. Notably, when this protein is inactive, there is a stark reduction in the activity of core anaerobic genes, underscoring the indispensability of MED25 for the full functionality of ERF-VII transcription factors (Fig. 6).

**Figure 6.**
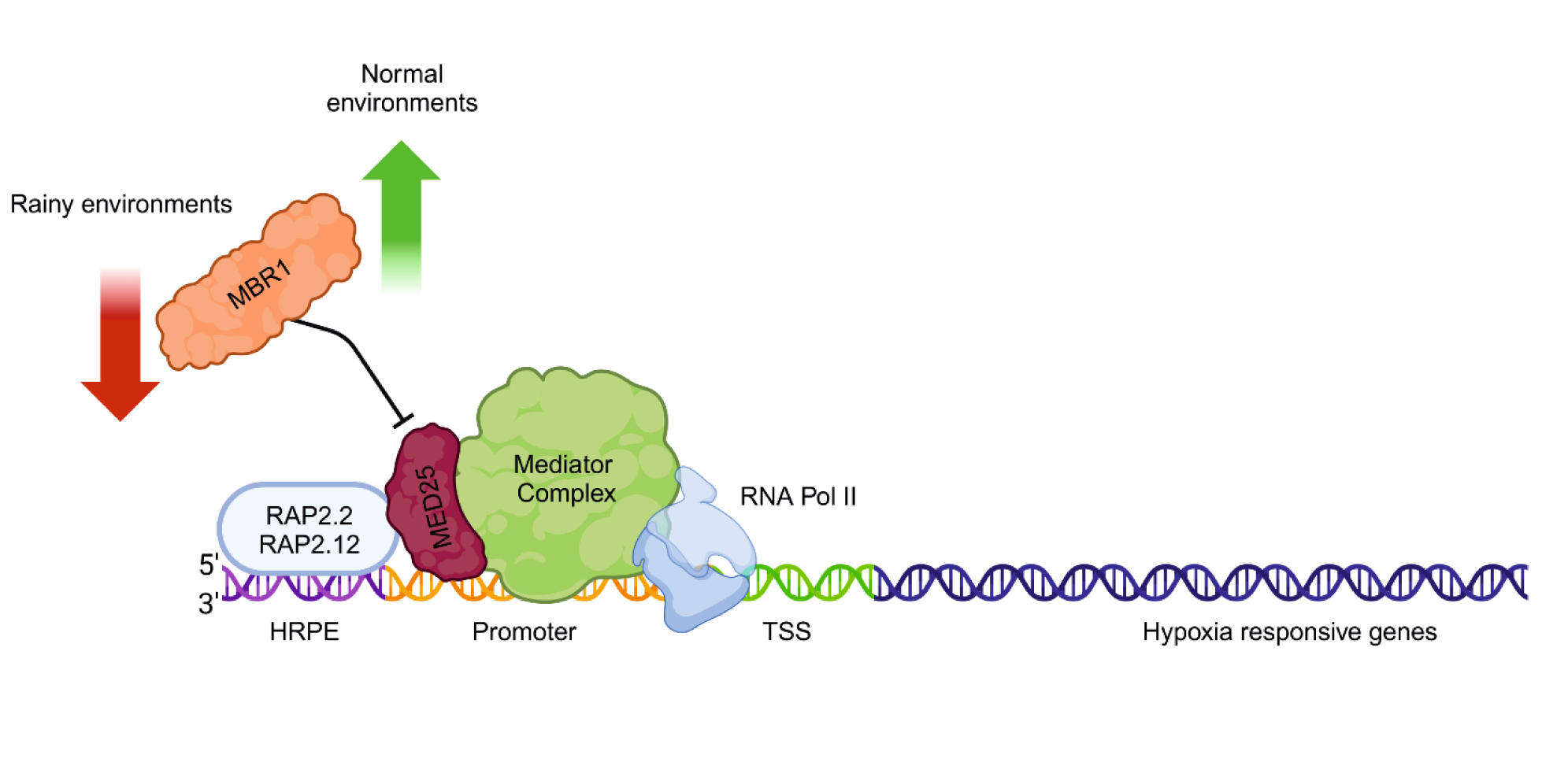
Proposed model based on our findings. Plants that inhabit particularly rainy environments have acquired mutations within the *MBR1* gene, leading to limited activity of MBR1 as a repressor of MED25. This limited activity results in an improved MED25 activity and, consequently, a more effective activation of the ERF-VII-dependent (RAP2.2; RAP2.12) hypoxic response in the plant.

In conclusion, our study aims to serve as a bridge connecting two domains of genetics: forward and reverse genetics, as well as an attempt to unite the potential of emerging tools, such as eGWAS, with cutting-edge molecular techniques. The divide between these two approaches, although providing valuable information independently, often leaves crucial gaps in our understanding of genetic mechanisms. By forging this connection, we aimed to paint a more comprehensive picture of the adaptation processes.

## Methods

### Plant materials and diversity analysis

The Arabidopsis accessions used in this study are part of the 1001 Genomes project ^60^, which consists of 1135 natural inbred *Arabidopsis thaliana* ecotypes (or accessions) with their respective DNA sequencing data. Starting with the overall dataset, accessions were filtered out to obtain the most informative set of ecotypes. Pairwise genome-wide identity-by-state (IBS) differences were calculated using PLINK v1.9 ^61^. When the pairs differed by less than < 0.01 changes per polymorphic site, we randomly removed one member of the pair. The VCF file with SNP calls from the 1001 Genomes project (https://1001genomes.org/data/GMI-MPI/releases/v3.1/) was therefore filtered with BCFtools v. 1.12 ^59^ resulting in 934 ecotypes retained (Supplemental Table 4).

The population structure and genetic clusters for the 934 Arabidopsis accessions were reconstructed using ADMIXTURE v. 1.3.0 ^63^. ADMIXTURE is a clustering software that infers population ancestries starting from highly informative, unlinked SNP data. The starting VCF file was restricted to biallelic SNPs with a genotype calling rate >95% and a minimum allele frequency >1% using BCFtools. The data set was then LD-pruned using PLINK. ADMIXTURE was then run with a 20-fold cross-validation (K=2 to K=20). The resulting best value of K, represented by the lowest cross-validation error value, was selected as the number of resulting clusters.

### Precipitation data, soil data and correlation analysis

Geospatial analyses were performed with the Geographic Information System software QGIS v3.16 ^64^. The GPS coordinates for each of the 934 accessions were derived from the 1001 Genomes Project database. The corresponding climatic data were retrieved from WorldClim (WC) ^62^. Rainfall measures were also derived from the climatologies at high resolution for the earth’s land surface areas (CHELSA) ^63^, the Climate Research Unit (CRU) ^64^ and the Global Precipitation Climatology Center (GPCC) ^65^. Association analyses for each individual month were performed using the WorldClim Historical monthly weather dataset^62^.

Considering the close correlation between hypoxic stress and soil water retention capacity, we extrapolated soil physical attributes from the SoilGrids 250m v2.0 database of the International Soil Reference and Information Centre (ISRIC) ^66^. Information on the bulk density, clay content, silt content, sand content and coarse fragments was collected. Full information on the pedoclimatic variables used in this study can be found in Supplemental Table 5.

Statistical analyses were conducted with R ^70^ custom scripts, unless stated otherwise. Precipitation and soil variables were checked for correlation using Pearson’s test (*r*). A PCA was performed to reduce the complexity and dimensionality of the rainfall dataset. To reveal the influence of precipitation patterns on the potential divergence of genetic clusters, we integrated precipitation data with the ADMIXTURE-based membership probabilities of each genome. Statistical analyses (Student’s *t*-test) were performed using R/*ggpubr* package, and plots were produced using R/*ggplot2* ^71^.

### eGWAS analysis

The eGWAS was conducted using an Efficient Mixed-Model Association eXpedited (EMMAX) ^69^ algorithm as implemented in the web-based application easyGWAS ^73^. EMMAX performs statistical tests for association mapping while also accounting for the population structure. EMMAX fits a model as: *Y* = *μ* + *X*_*i*_*β* + *Zu* + *ε*, where *Y* is the vector of trait values, *μ* is the mean value, *X* is the alternative allele dosage at SNP *i*, and *β* is the allelic effect of SNP *i* on the trait. The structure is corrected with a random genotype term represented by *u*, which follows a multivariate normal distribution *N*(0, *A*σ_G_^2^), where *A* is the relationship matrix between all individual genotypes built from the information of SNP information, and σ_G_^2^ is the genotype associated variance. SNPs were considered significantly associated with a pedoclimatic variable when they exceeded the Bonferroni threshold for a nominal test with *α* = 0.05. To further reduce the possibility of false positives, we focused on the genetic variants that showed the strongest statistical association (top hits). Quantile-Quantile (QQ) plots were utilized to assess the normality of the dataset and to identify any deviations from the expected distribution. The QQ plots were generated by plotting the quantiles of our sample data against the quantiles of a theoretical normal distribution (Fig. 1e; Supplemental Fig. 6 and 7). Manhattan plots and qqplots showing the results of association analyses were produced using R/*CMplot* ^74^. Linkage disequilibrium (LD) analysis was performed using PLINK v1.9 ^61^. LD analysis was performed on the genomic region containing SNPs with high significance located on Chr2 around the *MBR1* gene. The focus was confined to this specific region to ensure the identification and characterization of alleles that may be in strong LD with the significant SNPs. LD plot was generated using Haploview v4.2 ^75^.

### Plant material and growth condition

The experimental validation of the associations was performed on five accessions: Columbia (Col-0), serving as the wild-type reference genotype for all experiments, together with UKID96, UKID116, Ty-1, and Oy-0, which are natural inbred ecotypes belonging to the 1001 Genomes project. Inbred ecotypes were selected as they were sampled in regions with the highest rainfall. T-DNA insertion mutants for the *mbr1* and *med25* genes were also used. All seeds were obtained from the Nottingham Arabidopsis Stock Centre (NASC). The seeds were seeded in a soil mixture containing 70% professional potting medium and 30% perlite. They were then vernalized at 4°C in the dark for 48 hours. They were subsequently germinated at 22°Cday/18°C night, with a photoperiod of 12 hours at 120 *μ*mol photons m^−2^ s^−1^. The plants were grown in pots for 3-4 weeks before being used for the experiments.

### Submergence, Waterlogging and Phenotyping

To test plant responses to flooding and subsequent recovery from stress, plants were submerged (ZT6) in plastic tanks with the water level 10 cm above the leaf level and kept in the dark. For the comparison between the accessions and the Col-0 genotype, the plants were subjected to 72 h of submergence, then the plants were removed from the boxes and the photoperiodic conditions were restored (22°C day/18°C night with a photoperiod of 12 hours). Recovery from stress was evaluated one week after treatment ended. For the submergence experiment comparing Col-0, *mbr1* and *med25* knockout genotypes, the plats were subjected to 48h submergence, then the plants were removed from the boxes and the photoperiodic conditions were restored. Recovery from stress was evaluated after 48h after treatment ended. For waterlogging treatments, the root system of 3-week-old plants was immersed in water, but the leaves and petioles were left above the water level.

Before phenotyping, the plants were randomised to minimize potential bias and improve the reliability of our results by ensuring that variations in environmental conditions were evenly distributed. The resistance to stress was subsequently assessed under waterlogged conditions 20 days after the initiation of waterlogging. Resistance to stress was evaluated as the ratio of the leaf area of the control plants grown in air to the area of the plants that underwent submergence treatment, referred to as the Plant Leaf Area Ratio (PLA Ratio).

The plants were phenotyped using the LabScanalyzer (LemnaTec, GmbH, Aachen, Germany) digital phenotyping machine, equipped with a Manta G-1236 camera and a Kowa LM12XC lens. The plant trays were illuminated by two cool white LED panels mounted beside the camera at an angle of 30 ° to prevent direct reflection from the imaging area. The raw images were demosaiced using the AHD (Adaptive Homogeneity-Directed Demosaicing) algorithm from the OpenCV library and stored as 8-bit PNG images ^75^.

### RNA extraction and qPCR

Total RNA was extracted as previously described by Perata et al.^76^ with a minor modification (omission of aurintricarboxylic acid) to make the protocol compatible with the subsequent PCR procedures. Electrophoresis using 1% agarose gel was performed for all RNA samples to check for RNA integrity, followed by spectrophotometric quantification. Reverse transcription was performed using the Maxima First Strand complementary DNA (cDNA) synthesis kit for RT-qPCR, with dsDNase (Thermo Fisher Scientific). RT-qPCR was performed using 30 ng cDNA and iQ SYBR Green Supermix (BioRad Laboratories), according to the manufacturer’s instructions. *ACTIN2* expression (*ACT2*) was used as endogenous control for all genotypes analysed. A full list of the primers used for qPCR is provided in Supplemental Table 6.

### Construct preparation

For the preparation of the construct *35:MED25:FLuc*, the coding sequence (CDS) of the gene was amplified from cDNA from Col-0 plants. This was then cloned into the pENTR/D-TOPO vector, and the resulting entry vector was recombined into the plasmid *p2GW7L* using Gateway LR Clonase II (Thermo-Fisher Scientific). To generate the constructs *pMBR1:MBR1* and *pMBR1wet:MBR1wet,* the promoter and CDS of the gene were amplified from Col-0 and Ty-1 genotypes, respectively.

After recombination with *MBR1* CDS of *MBR1* and the variant version of the gene, the plasmid *p2GW7* was digested with *SacI* and *SpeI* to remove the CamV 35S promoter. The promoter of *MBR1* and the variant version were then ligated into the vector using a T4 Anza ligation mix (Thermo-Fisher Scientific). To produce the promoters of *PCO1*, *HYPOXIA RESPONSE ATTENUATOR1* (*HRA1*) and *ALCOHOL DEHYDROGENASE 1* (*ADH1*) fused to the firefly luciferase, the promoters of the respective genes was amplified starting with DNA extracted from WT plants (Col-0). The isolated promoters were then cloned into pENTR/D-TOPO and recombined in the destination vector *pGW7L*. Lastly, to produce overexpressors of *RAP2.2* and *RAP2.12*, the CDS of the genes were amplified from WT (Col-0) cDNA, cloned into pENTR/D-TOPO, and recombined in the destination vector. The list of all vectors used in this study is provided in Supplemental Table 7.

### Isolation and transformation

The protoplasts were isolated from leaves of 3-week-old plants by incubation in enzymatic solution (1% w/v cellulase, 0.3% w/v macerozyme, 0.4 M mannitol, 20 mM KCl, 10 mM CaCl_2_, 20 mM MES, pH 5.7) for 3 h in the dark at 22 ° C. Protoplasts were then filtered, washed twice with the W5 solution (154 mM NaCl, 125 mM CaCl_2_, 5 mM KCl, 2 mM MES, pH 5.7) and then centrifuged for 2 min at 100 × g before being resuspended in 0.4 M mannitol, 15 mM MgCl_2_, 4 mM MES (2-(N-morpholino)ethanesulfonic acid, pH 5.7) until a final concentration of 5 × 105 cells ml^−1^ was obtained. For transformation, 4 µg of each plasmid was added to 100 µl of protoplast suspension, and then gently mixed with an equal volume of a 40% PEG 4000 solution (0.2 M mannitol, 100 mM CaCl2). The mixture was incubated for 20 min at room temperature in the dark and then 440 µl of W5 solution were added to stop the transformation. The protoplasts were centrifuged at 100 × g. for 2 min, resuspended in 1 ml of 12 WI solution (50 mM mannitol, 4 mM MES pH 5.7, 20 mM KCl, 50 mM glucose) and transferred to six multiwell plates. The next day, protoplasts were pelleted by 3 min centrifugation at 5000 g, and flash frozen in liquid nitrogen for storage.

### Quantification of luciferase activity

The dual luciferase reporter assay system (Promega) was used to quantify the activities of Firefly (*Photinus pyralis*) and *Renilla reniformis* luciferase, according to the manufacturer’s instructions. In the case of protoplast transient transformation, firefly luciferase was normalized to Renilla luciferase activity using the Lumat LB 9507 tube Luminometer (Berthold).

## Author contributions

Conceptualization: SC, PP, EL, MD

Methodology: SC, PP, EL, MD

Investigation: SC, PMT

Supervision: PP, EL, MD

Writing—original draft: SC

Writing—review & editing: PP, EL, MD

**The authors declare no competing interests**

---

**Figure S1.**
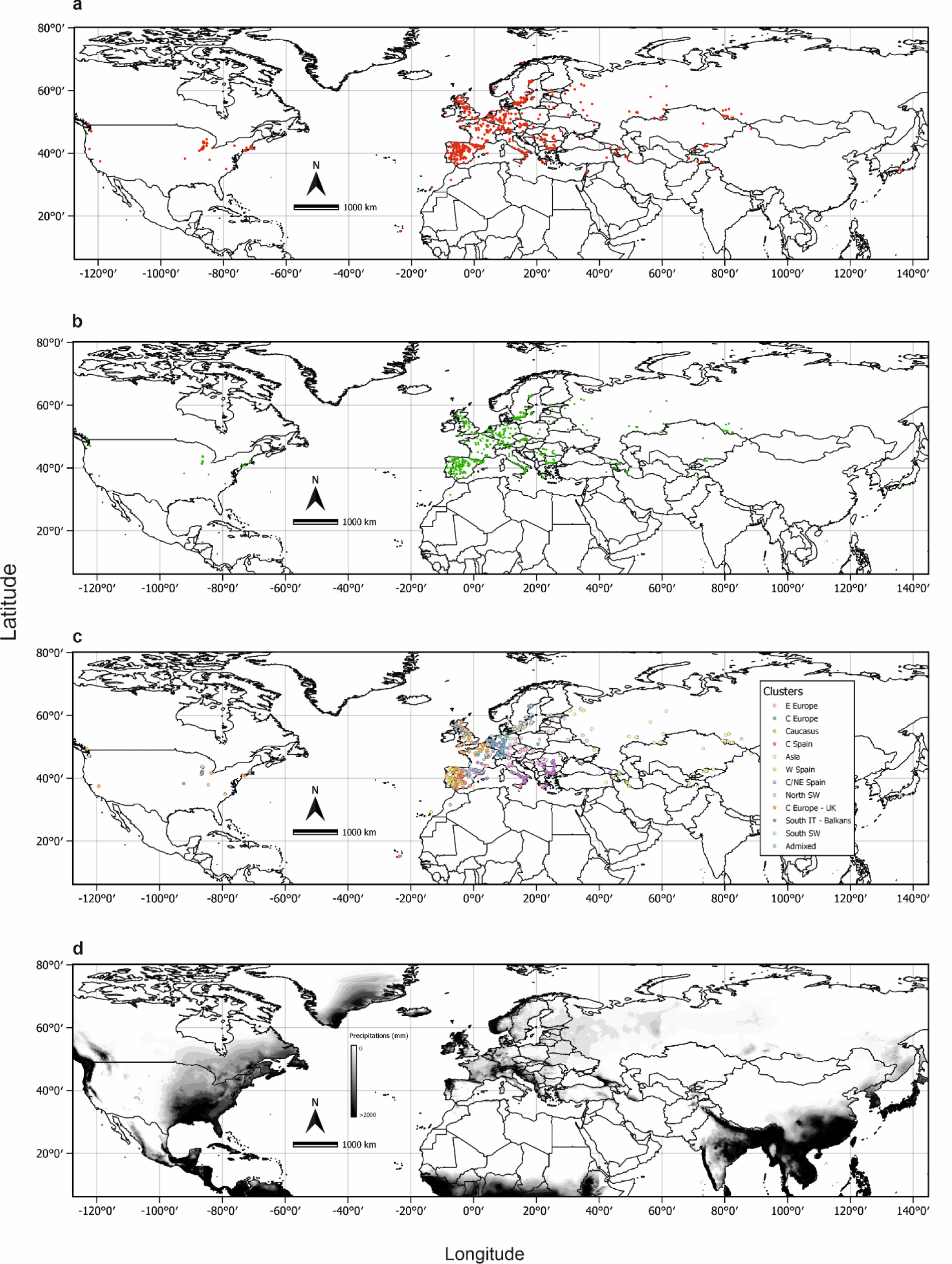
Maps representing: **(a)** the distribution of the complete dataset of the 1,335 accessions, **(b)** the distribution of the 934 accessions used for subsequent association analyses after filtering for IBS, **(c)** the distribution of the 11 clusters highlighted by the ADMIXTURE analyses, **(d)** the average annual rainfall for the period 1970 - 2020; the grey scale indicates the average rainfall expressed in millimetres of rainfall.

**Figure S2.**
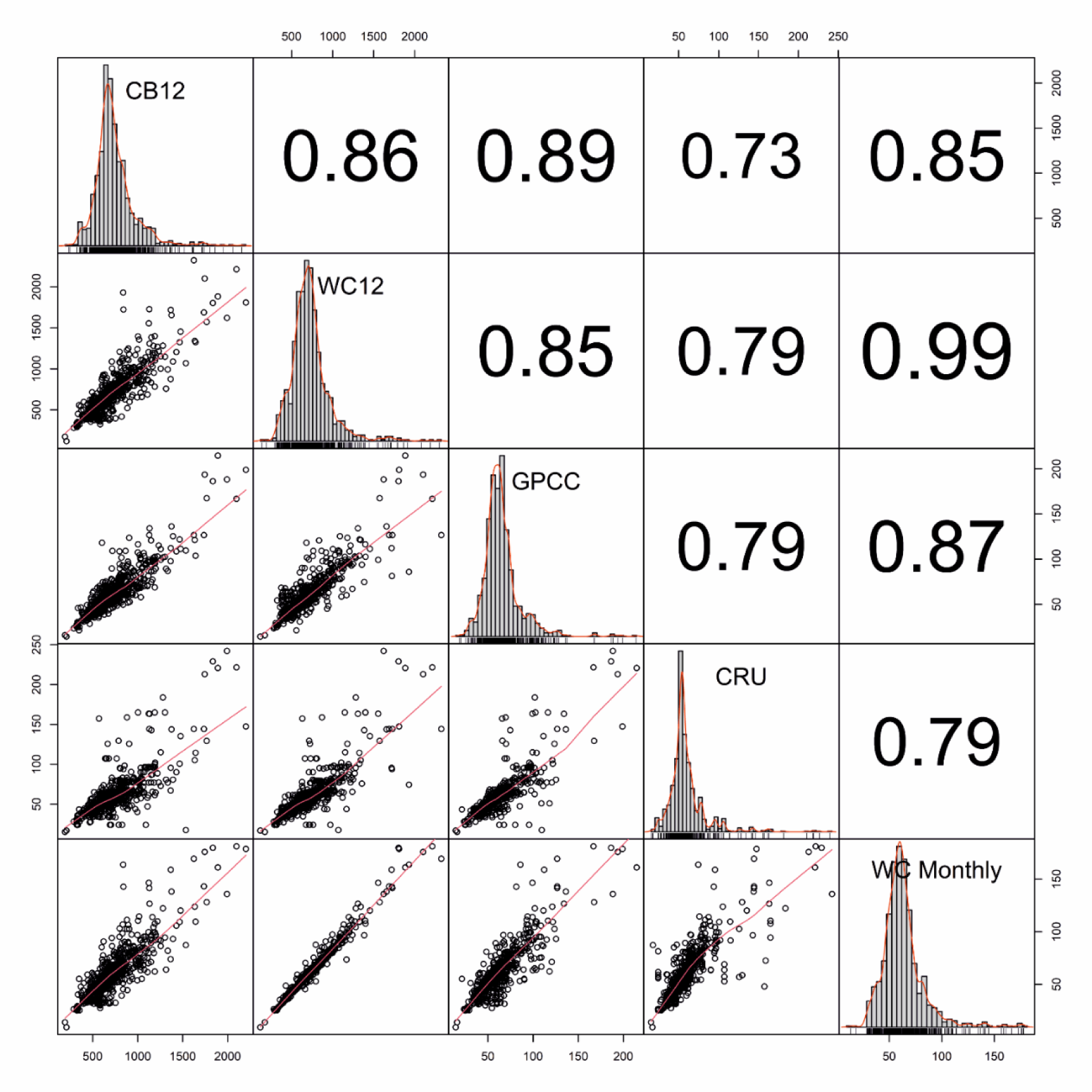
Correlation plot of the precipitation variables used in the study. Pearson’s correlation test (*r*) was used.

**Figure S3.**
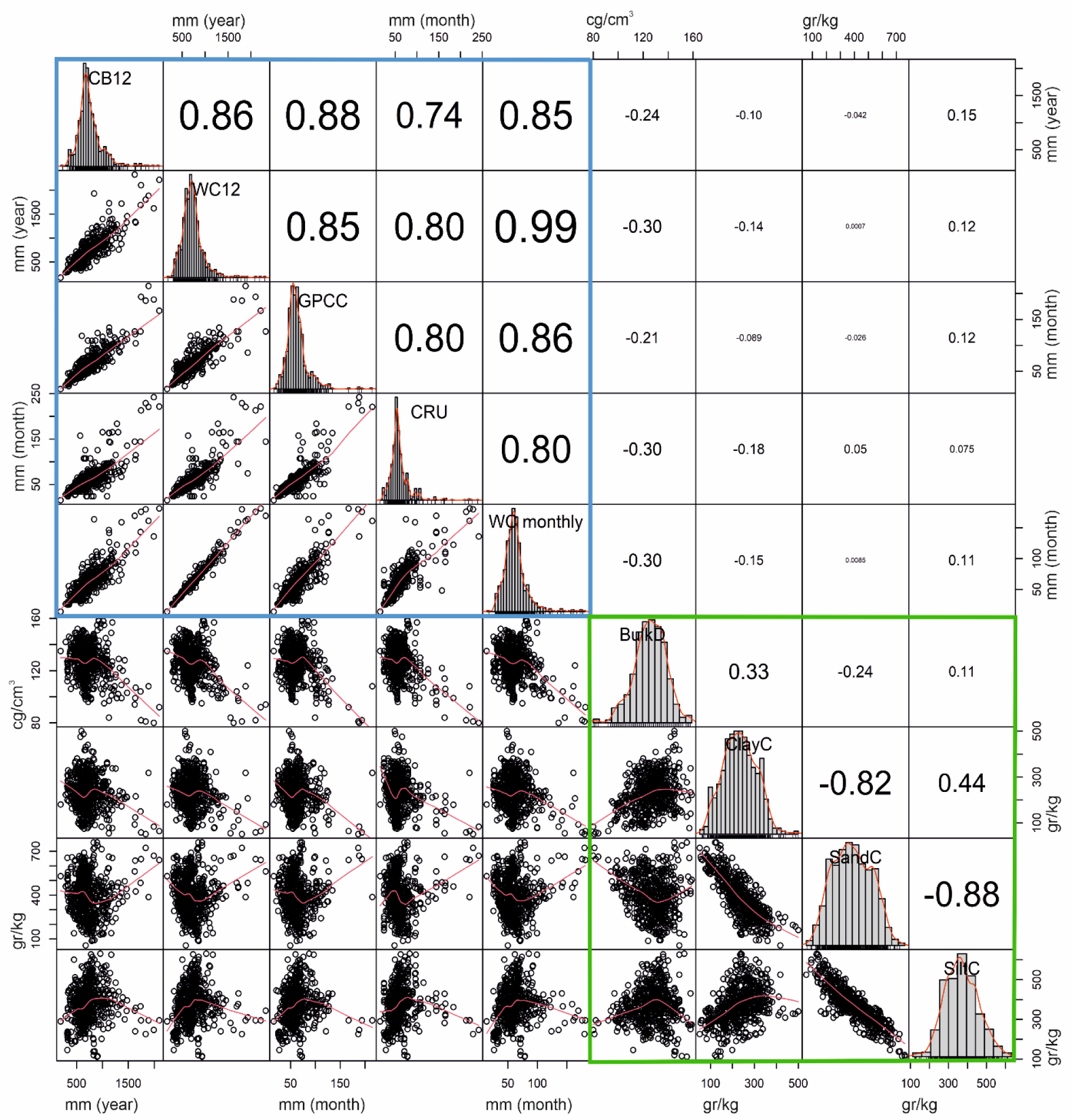
Correlation matrix for the precipitation and soil variables. The upper triangle contains Pearson correlation coefficients (*r*) between each pair of variables; the lower triangle displays scatter plots for the pairs. Diagonal elements show histograms representing the distribution of each variable. Precipitation variables are highlighted within the blue rectangle, while soil variables are highlighted within the green rectangle.

**Figure S4.**
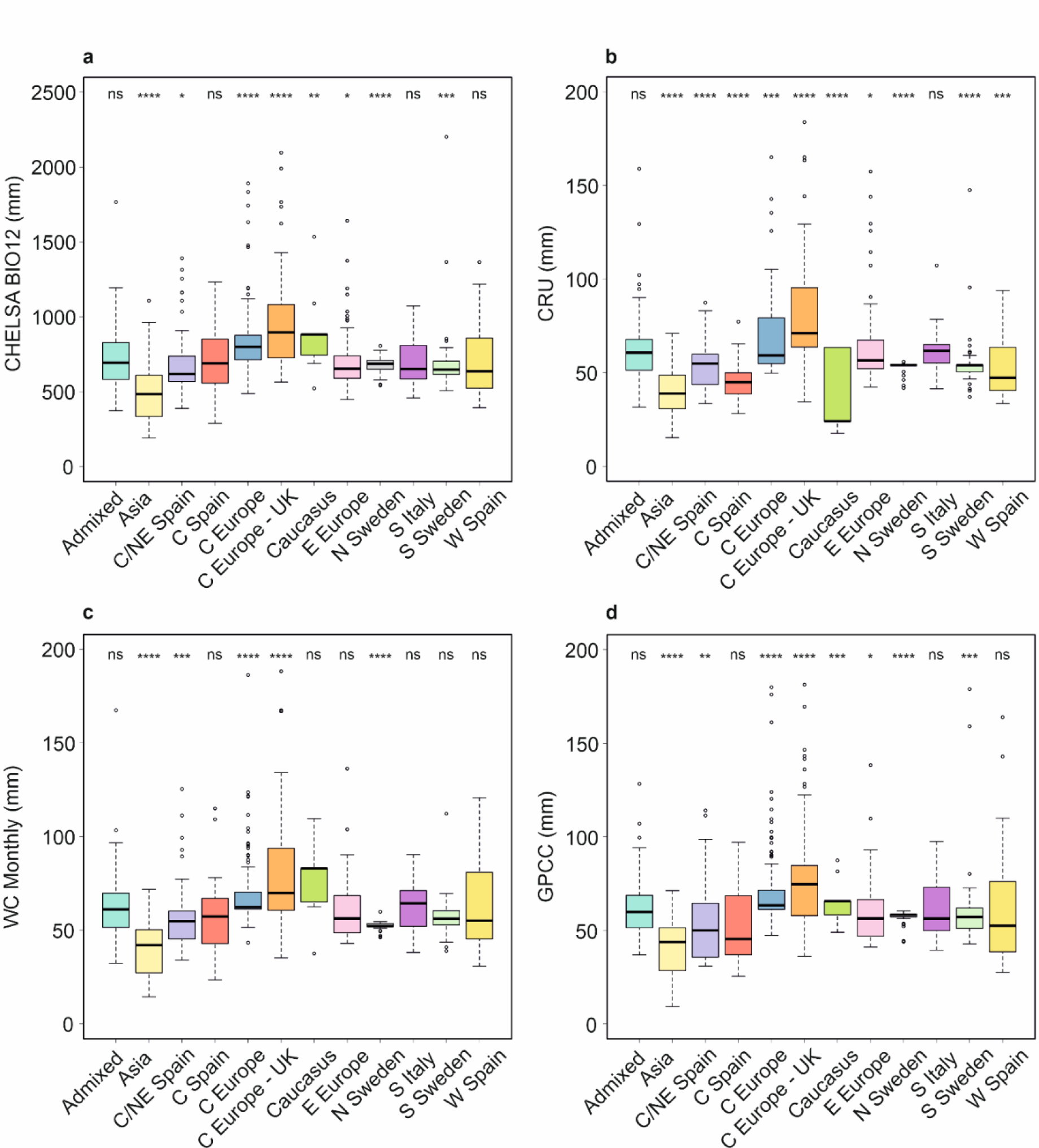
Distribution of the ecotypes based on the precipitation variables used in this study. The ecotypes are divided into the 11 clusters highlighted by the ADMIXTURE analyses; statistically significant differences are indicated by asterisks (Student’s t-test; ns = non-significant; * = p < 0.05; ** = p < 0.01; *** = p < 0.001; **** = p < 0.0001)

**Figure S5.**
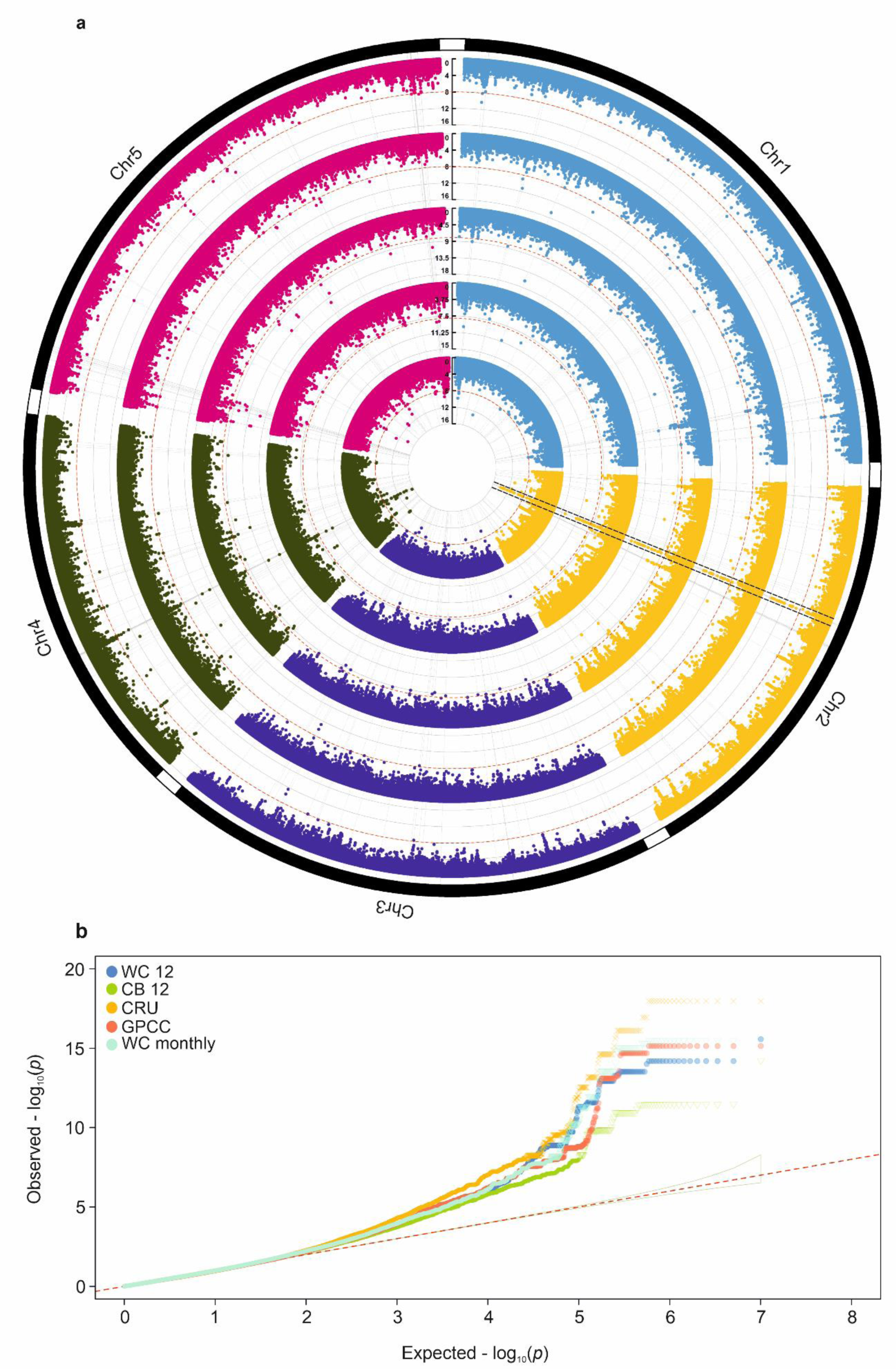
(a) Circular Manhattan plot for the eGWAS analyses of the rain variables. From outer to inner circle: WorldClim BIO 12; CHELSA BIO12; CRU; GPCC and WorldClim monthly precipitations; dashed lines indicate the position of the *MBR1* gene. **(b)** Multitrack Q-Q plot of the eGWAS analysis for the precipitation variables. The region outlined in light blue depicts the 95% confidence interval under the null hypothesis of a uniform *P* value distribution.

**Figure S6.**
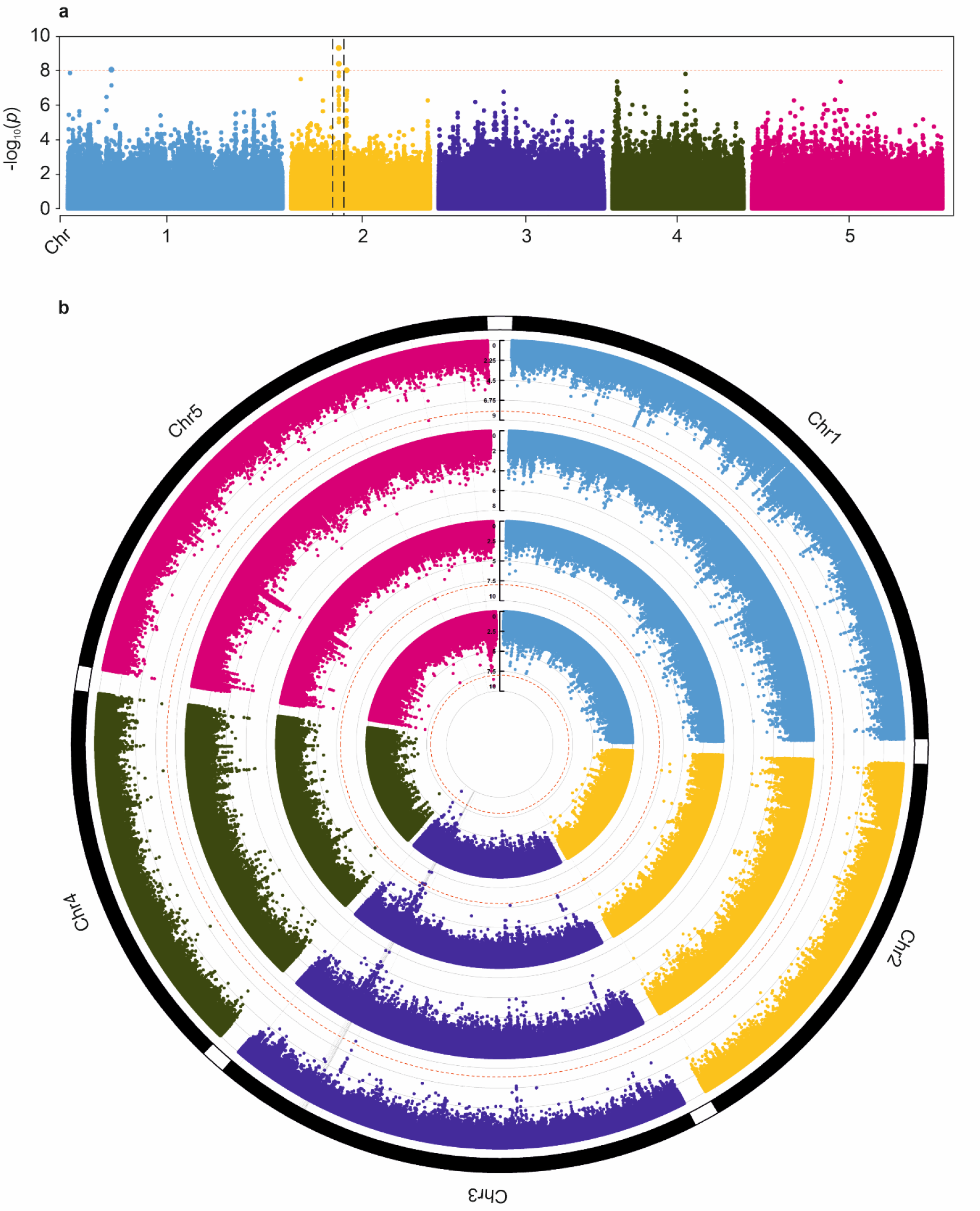
(a) Manhattan plot for bulk density; the dashed lines indicate the position of the *MBR1* gene. **(b)** Circular Manhattan plot for the pedological variables used in the study. From outer to inner circle: clay content; sand content; silt content; coarse fragments

**Figure S7.**
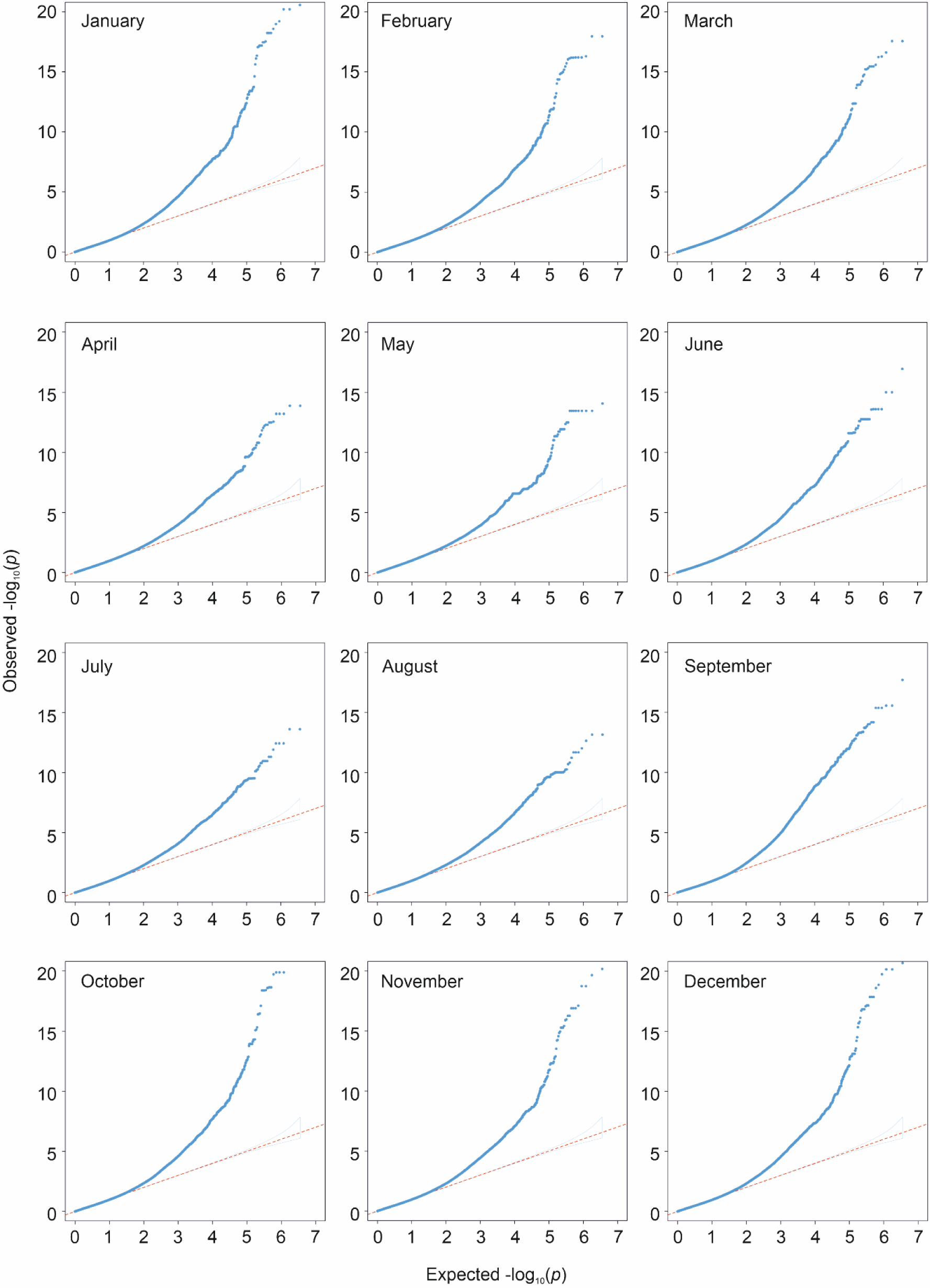
Q-Q plots of eGWAS for the monthly average precipitation data for the period 1901 – 2020; the regions outlined in light blue depict the 95% confidence interval under the null hypothesis of a uniform *P* value distribution.

**Figure S8.**
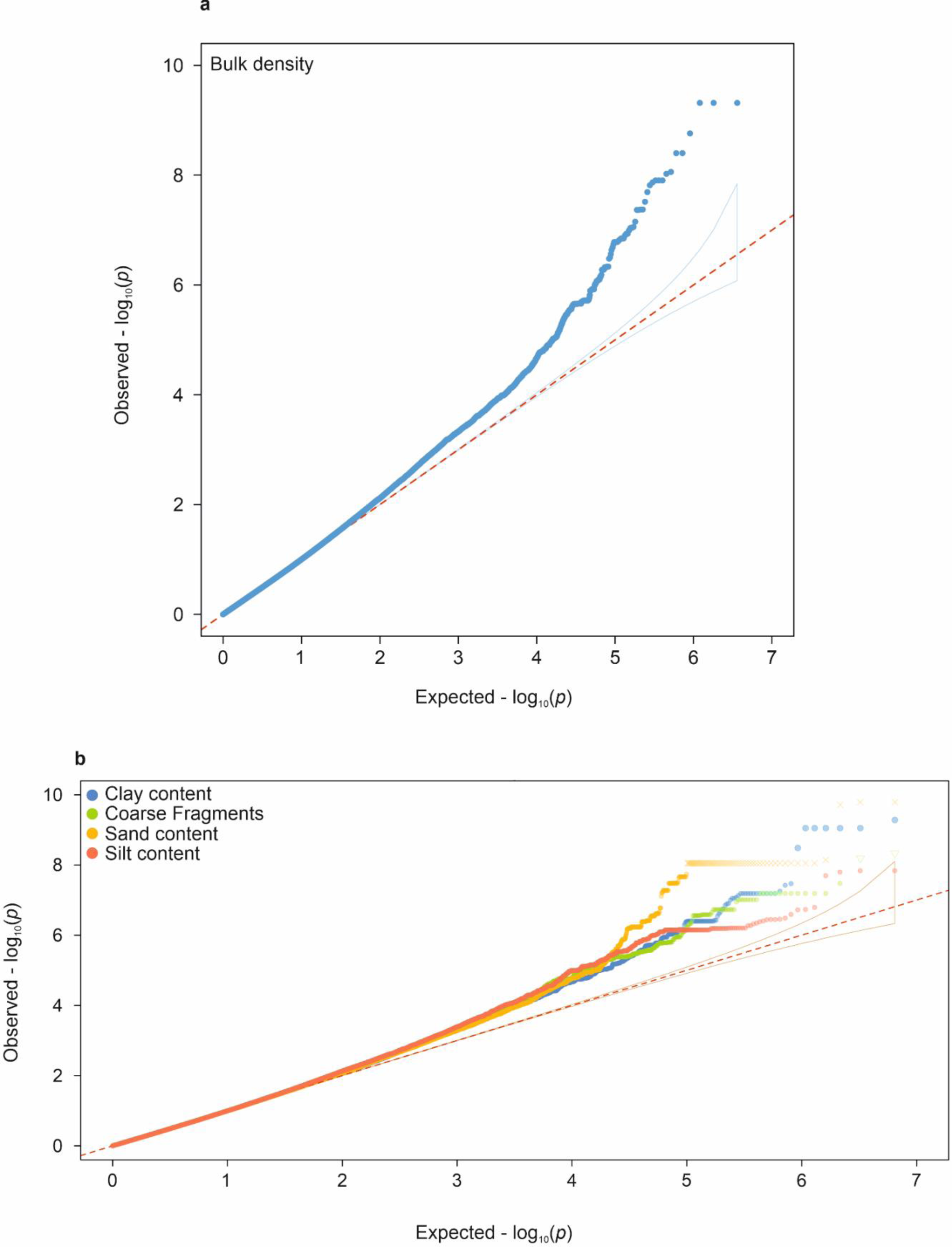
(a) Q-Q plots of eGWAS for bulk density. **(b)** Q-Q plots of eGWAS for soil variables; the regions outlined in light blue and light red depict the 95% confidence interval under the null hypothesis of a uniform *P* value distribution.

**Figure S9.**
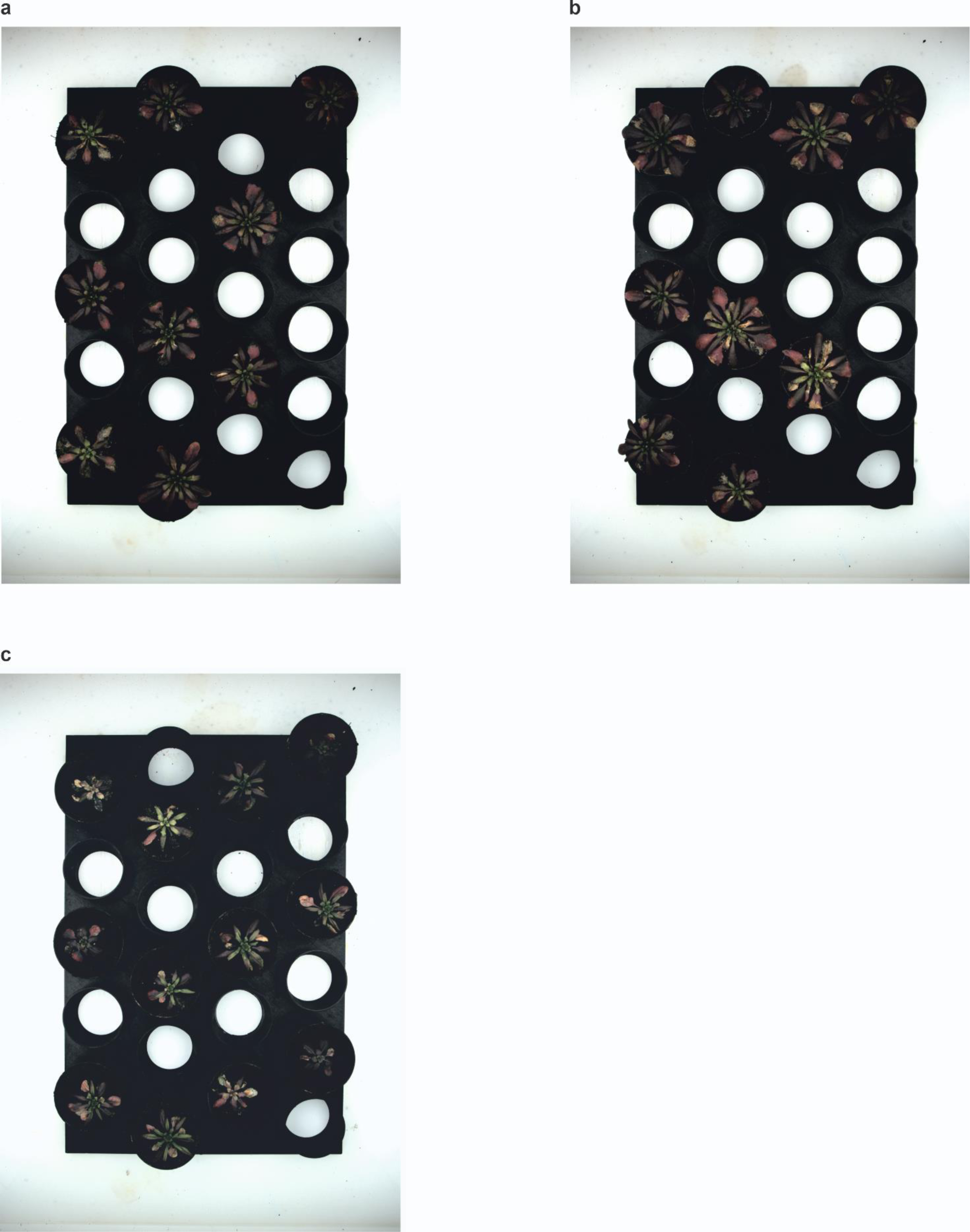
Unedited images sourced for Figure 2b (**a)** Col-0 (**b)** *mbr1* (**c)** *med25*

